# Contrast sensitivity reveals an oculomotor strategy for temporally encoding space

**DOI:** 10.1101/388132

**Authors:** Antonino Casile, Jonathan Victor, Michele Rucci

**Affiliations:** Istituto Italiano di Tecnologia, Center for Translational Neurophysiology, Ferrara (FE) 44121’Italy; Center for Neuroscience & Cognitive Systems, Rovereto (TN) 38068, Italy; Harvard Medical School, Department of Neurobiology, Boston, MA, 02115; Feil Family Brain and Mind Research Institute and Department of Neurology, Weill Cornell Medical College, New York, NY 10065, USA; Center for Visual Science, University of Rochester, Rochester, NY 14642, USA

**Author notes:** For correspondence (AC); (AC); (MR).

## Abstract

The contrast sensitivity function (CSF), how sensitivity varies with the spatial frequency of the stimulus, is a fundamental assessment of visual performance. The CSF is generally assumed to be determined by low-level sensory processes. However, the sensitivities of neurons in the early visual pathways, as measured in experiments with immobilized eyes, diverge from psychophysical CSF measurements in primates. Under natural viewing conditions, as in typical psychophysical measurements, humans continually move their eyes, drifting in a seemingly erratic manner even when looking at a fixed point. Here, we show that the resulting transformation of the visual scene into a spatiotemporal flow on the retina constitutes a processing stage that reconciles human CSF and the response characteristics of retinal ganglion cells under a broad range of conditions. Our findings suggest a fundamental integration between perception and action: eye movements work synergistically with the sensitivities of retinal neurons to encode spatial information.

## Introduction

Contrast sensitivity, the ability to distinguish a patterned input from a uniform background, is one of the most important measures of visual function ***(Robson, 1966; Campbell and Robson, 1968; De Valois et al., 1974; Owsley, 2003).*** Elucidation of its underlying mechanisms is, thus, essential for understanding how the visual system operates both in health and disease.

It has long been established that sensitivity varies in a specific manner with the spatial frequency of the stimulus, yielding the so-called contrast sensitivity function (henceforth CSF). Under photopic conditions, the CSF measured with stationary gratings exhibits a well-known band-pass shape that typically peaks around 3-5 cycles/deg and sharply declines at higher and lower spatial frequencies. The mechanisms responsible for this dependence on spatial frequency are not fully understood. At high spatial frequency, a decline in sensitivity is expected for several reasons, including the filtering of the eyes’ optics ***(Campbell and Green, 1965)*** and the spatial limits in sampling imposed by the cone mosaic on the retina ***(Hirsch and Miller, 1987; Rossi and Roorda, 2010).*** At low spatial frequencies, however, the reasons for a reduced sensitivity have remained less clear.

A popular theory directly links the low-frequency attenuation in visual sensitivity to the neural mechanisms of early visual encoding ***(Atick and Redlich, 1990, 1992).*** Building on theories of efficient coding ***(Barlow, 1961)***, it has been argued that this attenuation reflects a form of matching between the characteristics of the natural visual world and the response tuning of neurons in the retina: retinal ganglion cells (henceforth RGCs) respond less strongly at low spatial frequencies so as to counterbalance the spectral distribution of natural scenes. According to this proposal, this Altering eliminates part of the redundancy intrinsic in natural scenes and enables more efficient (***i.e.***, more compact) visual representations.

Although very influential, this proposal conflicts with experimental data. Neurophysiological recordings have long shown that the way the responses of retinal ganglion cells vary with spatial frequency deviates sharply from the CSF. The CSF of macaques is very similar to that of humans ***(De Valois et al., 1974);*** yet neurons in the macaque retina respond much more strongly at low spatial frequencies than one would expect from behavioral measurements of the CSF (Fig. 1A). This deviation cannot be reconciled with standard models of retinal ganglion cells. It persists even when one takes into account obvious differences in the stimuli often used in neurophysiological and behavioral measurements (***i.e.,*** drifting gratings vs. temporally modulated gratings), as well as the nonlinear attenuation in responsiveness at low spatial frequencies exhibited by some retinal ganglion cells ***(Derrington and Lennie, 1984; Croner and Kaplan, 1995; Benardete and Kaplan, 1997).*** This mismatch between neuronal and behavioral sensitivity indicates that additional mechanisms contribute to the CSF.

**Figure 1.**
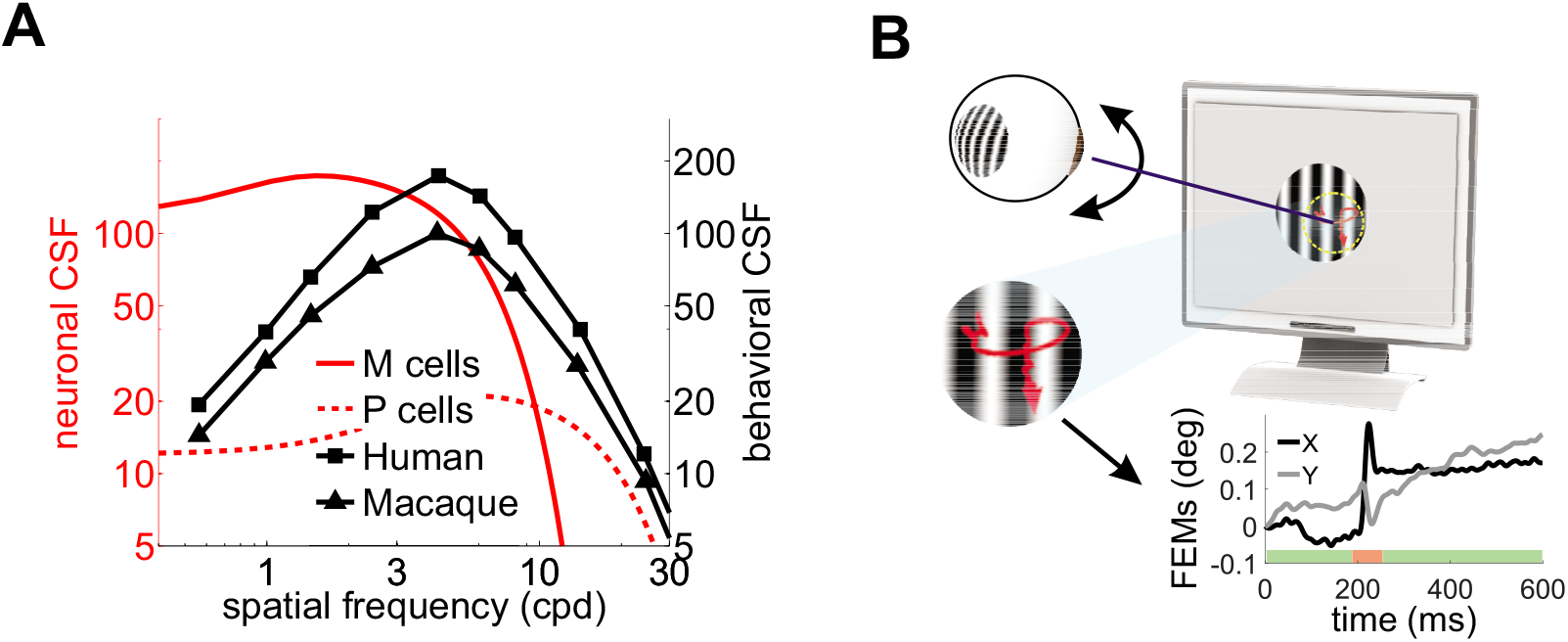
Contrast sensitivity and fixational eye movements. (A) Behavioral and neurophysiological measurements of contrast sensitivity. The contrast sensitivity functions (CSF) of humans and macaques (black curves;De ***Valois et al., 1974)*** are compared to the receptive fields profiles of magno-(M) and parvo-cellular (P) retinal ganglion cells (red curves; ***Croner and Kaplan, 1995).*** (B) Fixational eye movements (FEMs;red curve in magnified inset and black and gray traces in Cartesian graph), which include small saccades (microsaccades; red-shaded interval) and fixational drift (green), continually displace the stimulus on the retina.

A fundamental difference between neurophysiological and behavioral measurements of contrast sensitivity is the presence of eye movements in the latter. Under natural viewing conditions, humans and other primates incessantly move their eyes ***(Kowler, 2011; Cherici et al., 2012).*** Small movements, known as Axational eye movements (FEMs), occur, even when attempting to maintain steady gaze on a single point (Fig. 1 B). Although humans often tend to suppress saccades of all sizes, including microsaccades, during measurements of contrast sensitivity ***(Mostofi et al., 2016)***, ocular drift—the seemingly erratic motion in between saccades/microsaccades—keeps the stimulus on the retina always in motion and may cover an area as large as that of the foveola ***(Rucci and Poletti, 2015).*** Critically, this retinal image motion is completely eliminated or markedly attenuated in many neurophysiological preparations, where the retina is studied in a dish, or eye muscles are paralyzed as a result of anesthesia and/or neuromuscular blockade.

In previous work we have shown that eye drift profoundly reshapes the visual input signals to the retina, yielding temporal modulations that enhance high spatial frequencies ***(Casile and Rucci, 2009; Kuang et al., 2012; Aytekin et al., 2014).*** Humans are sensitive to these temporal signals and appear to use them for fine spatial discriminations ***(Rucci et al., 2007; Boi et al., 2017; Ratnam et al., 2017).*** These results have provided new support to the long-standing proposal that the visual system uses the luminance modulations generated from eye movements for encoding spatial information in a temporal format ***(Rucci and Victor, 2015).***

Building upon this previous work, here we investigate whether this encoding strategy, coupled with the known response characteristics of retinal neurons, accounts for the most fundamental properties of human spatial sensitivity. In addition to the properties described above, it is well established that contrast sensitivity is affected by temporal modulations in the stimulus. Whereas the CSF exhibits a strong attenuation at low spatial frequencies when tested with stationary gratings, the shape of this function changes when gratings are modulated in time, transitioning from bandpass to low-pass as the temporal frequency of the stimulus increases ***(Robson, 1966).*** Furthermore, although strongly attenuated, sensitivity also tends to shift to higher spatial frequencies when retinal image motion is strongly reduced, as in experiments of retinal stabilization ***(Kelly, 1979).*** In both these conditions, the temporal modulations impinging onto retinal receptors differ drastically from those generated by normal eye drift over stationary gratings.

Does a temporal strategy of spatial encoding reconcile neurophysiological and behavioral measurements of contrast sensitivity? And does consideration of the temporal structure of the retinal input explain the changes in the CSF measured in different experimental conditions? More broadly, does the oculomotor-driven dynamics of retinal ganglion cells provide a unifed account of human spatial sensitivity? Answers to these questions are not only critical for advancing our comprehension of the mechanisms of visual encoding but also for understanding the consequences of abnormal retinal image motion and their clinical implications.

In the following, we use neuronal models to quantitatively examine the impact of the fixational motion of the retinal image and compare the responses of retinal ganglion cells to the CSF of primates.

## Results

Fig. 1A compares the mean receptive fields of ganglion cells in the primate retina, as estimated by ***Croner and Kaplan (1995)***, with the contrast sensitivity of alert and behaving macaques ***(De Valois et al., 1974).*** The two sets of data deviate considerably, especially at low spatial frequencies. In this range, unlike the CSF, neural sensitivity is not strongly attenuated, a trend reported by multiple neurophysiological studies ***(e.g., Kaplan and Shapley, 1982; Hicks et al., 1983; Derrington and Lennie, 1984).*** This cannot be the result of an extrapolation of the receptive-field measurements, which were made at spatial frequencies down to 0.07 cpd ***(Croner and Kaplan, 1995).***

While a difference-of-Gaussians model can yield reduced responses at low spatial frequencies, attenuation similar to that observed in the CSF can only be achieved at the expense of highly unrealistic model parameters. As shown in Figs. S1A-B, for both M and P cells, matching the physiological CSF requires a surround strength that is more than twice the value found in physiological measurements, a condition that gives an almost perfect balance between excitation and inhibition. Even small deviations from this balance lead to marked departures from the CSF (Fig. S1A-D). Thus, the spatial sensitivity of retinal ganglion cells, considered in isolation, appears quantitatively incompatible with the idea that the CSF is determined by the spatial sensitivity of retinal neurons ***(Atick and Redlich, 1990, 1992).*** This proposal requires a greater attenuation of neural sensitivity at low spatial frequencies to counterbalance the large power of natural scenes in this range.

The response of a neuron, however, does not only depend on the cell spatial preference but also on its temporal sensitivity. Temporal transients are always present in the input signals to the retina during behavioral measurements of contrast sensitivity. Experimenters often take great care to minimize unwanted sources of temporal modulations, ***e.g.,*** by slowly ramping up the stimulus at the beginning and down at the end of a trial and by enforcing fixation to prevent visual transients caused by saccadic eye movements (Fig. 2A). Yet, despite these precautions, fixational eye movements are always present and modulate the visual flow impinging on the retina even when the stimulus does not change on the monitor. Could these oculomotor fluctuations result in a transformation that reconciles neurophysiological and behavioral measurements of spatial sensitivity?

**Figure 2.**
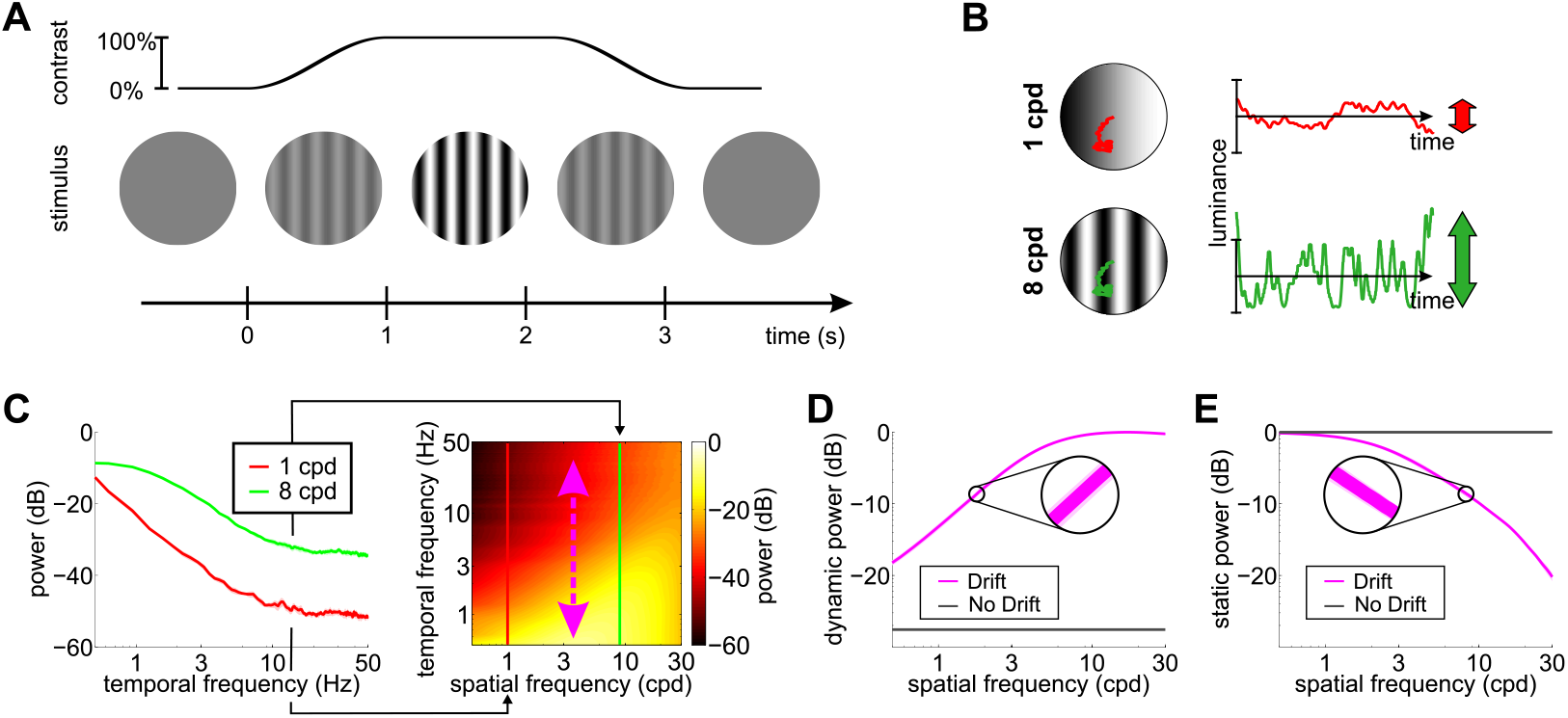
Input transients during measurement of contrast sensitivity. (A) Temporal modulations in the stimulus. Measurements of contrast sensitivity often change gradually the contrast of the stimulus during the course of the trial. In this case, the stimulus is a static grating. (B) Fixational jitter modulates input signals, even with a static stimulus. The amplitude of these modulations tend to increase with the spatial frequency of the grating (arrows). That is, the same fixational drift will produce bigger temporal modulations of the retinal input in the presence of gratings with higher spatial frequency ***(Rucci and Victor, 2015).*** (C) Temporal power distribution of the retinal inputs with gratings at 1 and 8 cycles/deg (left panel). Higher spatial frequencies lead to broader temporal distributions (right panel). (D) Dynamic power resulting from ocular drift as a function of spatial frequency (Drift). Data represent the total power integrated across non-zero temporal frequencies (purple vertical arrow in C) averaged over N = 5 observers. (E) The power remaining on the 0 Hz axis. In both D and E, the shaded regions represents one standard deviation (see insets). The power given by the same stimuli in the absence of eye movements, but taking into account the temporal envelope of the onset and offset of the stimulus, is also shown (No Drift).

To investigate this question, we recorded eye movements in human observers, as they carried out a grating detection task at threshold and exposed spatiotemporal Alters approximating the receptive f elds of retinal ganglion cells to the luminance signals experienced by the retina in each individual trial. Fig. 2B shows the temporal modulations impinging onto retinal neurons during a typical measurement of contrast sensitivity. In the absence of any transient, the power of a stationary visual stimulus would be conf ned to the DC (0 Hz) temporal frequency axis. In practice, however, both eye drift and the turning of the stimulus on and off on the display introduce temporal modulations. These modulations effectively redistribute part of the stimulus DC power to nonzero temporal frequencies, ***i.e.*** they transform static power (the original power at 0 Hz) into dynamic power (power at non-zero temporal frequencies).

As shown in Figs. 2C-D, because of the characteristics of ocular drift, the resulting dynamic power increases with spatial frequency, up to approximately 30 cpd, which, interestingly, roughly corresponds to the frequency limit given by the spatial resolution of photoreceptors in the fovea. In contrast, unlike drift, contrast modulations due to the onset/offset of the stimulus on the display cause power redistributions that do not depend on the spatial frequency of the stimulus (gray lines in Fig. 2 D-E). It is important to keep in mind that eye movements do not generate new power in the retinal input. They only redistribute the original DC power of the stimulus, so that a complementary frequency-dependent attenuation of power occurs along the 0 Hz axis (Fig. 2E).

Both eye drift and contrast changes yield temporal modulations that are well within the range of temporal sensitivity of retinal ganglion cells. However, in simulations that replicated the standard conditions of contrast sensitivity measurements, drift modulations predominated. Since drift modulations convey little power at low spatial frequencies, the responses of standard ganglion cells were attenuated in this frequency range (Fig. S2A). This happened for both M and P cells, despite the well-known differences in their spatio-temporal sensitivity. As a consequence of this effect, a simple linear combination of the resulting M and P responses accurately predicted human contrast sensitivity with stationary stimuli over the entire range of relevant spatial frequencies (solid line in Fig. 3).

**Figure 3.**
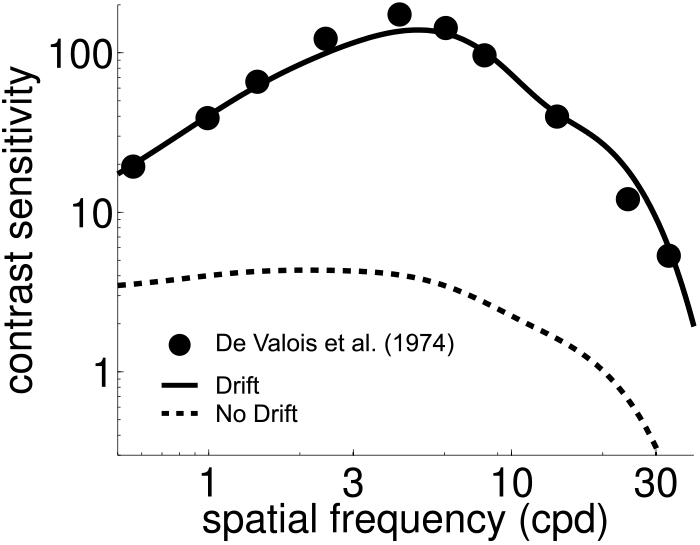
Influence of fixational drift on contrast sensitivity. Predicted CSFs in the presence (Drift; solid line) and absence (No Drift; dashed line) of eye movements. Stimuli were stationary gratings. A linear combination of the responses of M and P cells closely matches classical measurements (circles; data from ***De Valois et al., 1974***) only when eye drift occurs. CSFs predicted separately from the responses of M and P neurons are shown in Fig. S2 A.

In contrast, in the absence of eye movements, when the only temporal modulations were those given by the onset/offset of the stimulus on the monitor, the CSF predicted by the same linear combination of neural responses exhibited a low-pass behavior that deviated considerably from human contrast sensitivity, especially at low spatial frequencies (dashed line in Fig. 3). In fact, no linear combination of modeled responses could approximate the CSF in this condition. This happened because, unlike the luminance modulations resulting from ocular drift, the amplitude of the contrast modulations of the stimulus on the display does not depend on the spatial frequency of the stimulus (Fig. 2D). Thus, without taking ocular drift into account, neuronal models exhibit a higher level of response at low spatial frequencies, as dictated by the spatial sensitivity of their kernels — and this strongly deviates from the CSF (Fig. 1A). In sum, standard models of the responses of M and P RGCs well predict the shape of the human CSF as measured with stationary gratings, but only when the consequences of f xational drift on the retinal input are taken into account.

Contrast sensitivity is a function not only of the spatial frequency of the stimulus but also of its temporal frequency. Measurements with gratings modulated in time have long shown that the CSF in humans is not space-time separable: the way contrast sensitivity varies with spatial frequency depends on the temporal frequency of the modulation ***(Robson, 1966).*** As the temporal frequency increases, the CSF changes its shape, transitioning from band-pass to low-pass (Fig. 4A).

**Figure 4.**
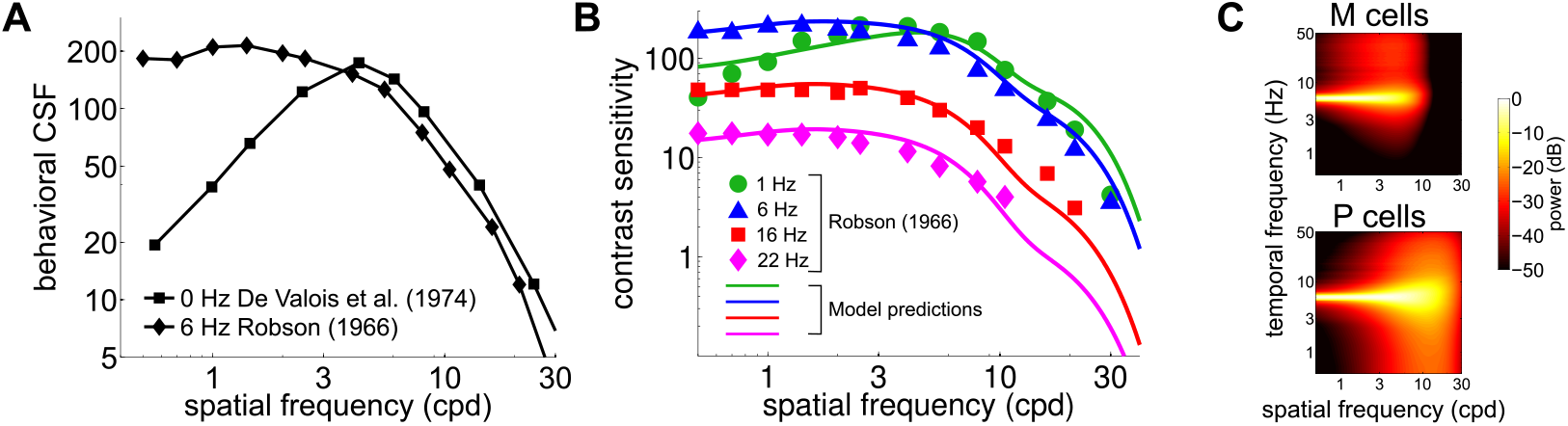
Contributions of fixational drift to contrast sensitivity with temporally modulated gratings. (A) Human CSFs measured with static (0Hz) and sinusoidally modulated (6Hz) gratings. Data at 0Hz are replotted from Fig. 1 A; 6Hz data are from ***Robson (1966).*** (B) Contrast sensitivity functions predicted by our model in the presence of temporally modulated gratings. Data points are measurements from ***Robson (1966)***. CSFs predicted separately from the responses of M and P neurons are shown in Fig. S1 B. (C) Power spectra of the response of modeled retinal ganglion cells during viewing of gratings temporally modulated at 6Hz. Each point in the map represents the amount of power at a given temporal frequency resulting from translating the modeled receptive fields over a grating at the corresponding spatial frequency following the recorded eye drift trajectories.

To investigate whether our model also accounts for this change in shape, we repeated our simulations using gratings modulated at various temporal frequencies. The same linear combination of the responses of M and P cells as in Fig. 3 continued to closely match human performance when the stimulus was temporally modulated on the display, and the predicted CSF replicated the low-pass to band-pass transition observed in primates, as the frequency of the modulation increased (Fig. 4B).

This change in shape was the consequence of the different amount of dynamic power that the combination of fixational drift and temporal modulations of the stimulus delivered within the range of neuronal sensitivity. Since our model assumes that there is no sensitivity to unchanging stimuli, the DC power does not contribute to cells’ responses. However, flickering a grating has the effect of shifting the 0Hz power to the temporal frequency of the modulation (Fig. 4C). As a consequence, as the frequency of the modulation increased, this DC power was progressively moved into the sensitivity range of modeled neurons. At low temporal modulating frequencies (e.g., 1Hz or below), only a small fraction of this power was within the region of neuronal sensitivity, and the temporal redistribution resulting from eye drift continued to exert a strong influence, forcing the CSF to maintain its band-pass shape. However, at higher temporal frequencies (e.g., 6Hz and higher), the power restricted to the 0Hz axis in the absence of stimulus’ modulations now became fully available within the cells’ peak sensitivity region. Since this static power is predominantly at low spatial frequencies (Fig. 2E), it caused a transition from band-pass to low-pass behavior in the responses of simulated M and P neurons, as well as in the shape of the CSF.

In sum, our model attributes changes in the CSF to the structure of temporal power that eye drift and contrast modulations in the stimulus deliver within the range of sensitivity of retinal ganglion cells. The model explains not only contrast sensitivity with stationary gratings, but also the bandpass to low-pass transition that occurs with temporally modulated gratings. Notably, it correctly predicts the temporal frequency range at which this transition takes place. Our results, thus, suggest a functional link between the physiological instability of visual fixation and the characteristics of the CSF.

A natural question then emerges: how is contrast sensitivity affected by elimination of the luminance modulations caused by ocular drift? Ideally, in the complete absence of eye movements, neural responses in our model would only be driven by the modulations present in the external stimulus. Under such conditions, the model predicts that sensitivity to a stationary grating would be greatly attenuated and the CSF would shift toward a low-pass shape, as it would lack the frequency-dependent amplif cation operated by ocular drift.

In real experiments, however, elimination of oculomotor-induced luminance modulations is impossible. Retinal stabilization — a laboratory procedure that attempts to immobilize an image on the retina ***(Riggs et al., 1953; Yarbus, 1957)*** — is always affected by noise in the oculomotor recordings as well as imperfections in gaze-contingent display control, which leave some residual motion on the retina. Under these conditions, contrast sensitivity has indeed been found to be attenuated but it maintains its band-pass shape and peaks at higher spatial frequencies ***(Kelly, 1979).***

To examine whether sensitivity to temporal transients accounts for the changes in the CSF measured under retinal stabilization, we exposed modeled neurons to reconstructions of the visual input signals experienced in these experiments. Previous studies have established that a Brownian model well captures the characteristics of retinal image motion during fixation ***(Kuang et al., 2012; Poletti et al., 2015).*** Building on this previous finding, we modeled the residual motion of the retinal image in stabilization experiments as a Brownian process, but with greatly reduced diffusion coefficient relative to that present during normal, unstabilized fixation.

Fig. 5A shows how the spatial frequency content of the luminance fluctuations experienced by retinal receptors (the power available at nonzero temporal frequencies) varies with the scale of the Brownian motion process ***(i.e.,*** its diffusion coefficient, D). Changing the amount of retinal image motion has interesting repercussions on the characteristics of temporal modulations. As expected, a smaller diffusion constant delivers less dynamic power to the retina within the range of neural sensitivity, a direct consequence of the fact that luminance modulations are now smaller. However, a smaller ***D*** also has the effect of shifting the range of amplification to higher spatial frequencies by a factor of 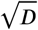. This happens because reducing the scale of retinal image motion is functionally equivalent to spatially stretching the stimulus, thus altering its frequency content.

These effects in the spectral distributions of the retinal flow well match the changes in contrast sensitivity observed in retinal stabilization experiments. Fig. 5B compares classical retinal stabilization data from ***Kelly (1979)*** to the sensitivity predicted by our model when the diffusion constant of the retinal image motion was attenuated by a factor of 125, which corresponds to shrinking the spatial scale of eye movements by approximately one order of magnitude. Model predictions closely followed psychophysical measurements: a reduction in the amount of retinal image motion attenuated contrast sensitivity while maintaining its band-pass shape and shifted its peak sensitivity to higher spatial frequencies from 4Hz to 5.5Hz (Fig. 5B).

These data show that consideration of the luminance modulations resulting from the motion of the stimulus on the retina accounts not only for behavioral sensitivity measurements performed in the presence of normal eye movements, but also for measurements made under conditions of retinal stabilization, when retinal image motion is greatly reduced.

## Discussion

Contrast sensitivity is a fundamental descriptor of visual functions. In many species, including humans, sensitivity strongly depends on the spatial and temporal frequency of the stimulus. Here we show that a temporal scheme of spatial encoding, a scheme in which spatial vision is primarily driven by temporal changes, predicts such dependencies when the temporal modulations introduced by incessant eye movements are taken into account. In contrast, when these consequences of fixational drift are ignored, the known response characteristics of retinal ganglion cells fail to account for human CSF. As described below, these results are highly robust, bear multiple consequences, and lead to important predictions.

An important consequence of our results regards the strategies by which the visual system encodes spatial information. Existing theories of visual processing have attributed the shape of the CSF to the characteristics of early visual processing. In an influential study, ***Atick and Redlich (1992)*** found that the theoretical filter that optimally decorrelates natural images closely matches the CSF. Since decorrelated responses enable compact neural representations, these authors assumed that the CSF reflects the average spatial selectivity of ganglion cells in the retina. However, experimental measurements have long shown that the response selectivity of RGCs differs considerably from the CSF, particularly at low spatial frequencies, where decorrelation would be most beneficial ***(Hicks et al., 1983; Kaplan and Shapley, 1982; Derrington and Lennie, 1984; Croner and Kaplan, 1995).*** As expected from this deviation, broad spatial correlations in RGCs responses have been found in preparations in which natural images are displayed in the absence of eye movements ***(Puchalla et al., 2005; Segal et al., 2015).*** These findings are consistent with our model: when the transients in stimulus presentation override the consequences of eye drift, spatial sensitivity follows the spatial kernels of modeled receptive fields. For this reason, responses to low spatial frequencies are enhanced relative to the level that would be needed for decorrelating activity.

The same principle also provides an explanation for the band-pass to low-pass transition of the CSF as the temporal frequency of the stimulus increases. This transition is the consequence of the dynamic power that the combination of fixational drift and stimulus transients delivers within the range of neuronal temporal sensitivity. With stationary gratings, temporal modulations in the retinal input are heavily influenced by ocular drift, which enhances high spatial frequencies imposing a band-pass sensitivity (Fig. 3). With temporally modulated gratings, neuronal responses are also affected by the contrast modulation imposed to the stimulus on the display. Above a frequency of a few Hz, the impact of external modulations outweighs the effects of eye movements, removes the space-time inseparability in cell responses caused by ocular drift, and enhances again sensitivity to low spatial frequencies (Fig. 4 B).

More broadly, our model relies on the fundamental assumption that the visual system encodes spatial information by means of the oculomotor-driven dynamics of neural responses. According to this proposal, the visual system is minimally sensitive to stimuli confined to 0Hz and relies on temporal changes for encoding space. During natural viewing, eye movements are a major source of temporal modulations to the retina, and the spatial information conveyed by these modulations critically depends on the way the eyes move. Thus, rather than attributing spatial sensitivity solely to the spatial selectivity of RGCs, our analysis shows that the CSF is shaped by the joint ***spatial*** and ***temporal*** characteristics of retinal responses and how they interact with oculomotor transients. It is striking that standard models of RGCs predict the CSF so well when exposed to the temporal modulations present in the experiments. While our study cannot exclude that other mechanisms, at various stages of visual processing, may also play a role in shaping the CSF (e.g., the number of neurons in different frequency channels), it suggests that these other contributions are minimal. Consideration of RGCs temporal sensitivity provides a parsimonious unifying framework for a wide range of experimental measurements of the CSF with only a minimal set of assumptions.

We specifically focused on fixational drift both because of its ubiquitous presence and its known influence on fine pattern vision ***(Ratliff and Riggs, 1950; Ditchburn, 1955; Steinman et al., 1973; Rucci et al., 2007; Ratnam et al., 2017).*** Other types of eye movements, like saccades and microsaccades, tend to be suppressed during measurements of contrast sensitivity ***(Mostofi et al., 2016)*** and were not considered in this study. The transients from these movements, however, differ in their spectra from those from eye drift, as they provide equal temporal power across a broad range of spatial frequencies. Thus, during normal viewing, the visual system could benefit from different types of modulations. In keeping with this idea, it has been argued that the stereotypical alternation of oculomotor transients resulting from the natural saccade/drift cycle contributes to a coarse-to-fine processing dynamics at each visual fixation ***(Boi et al., 2017).***

It is worth emphasizing that our results are very robust and do not depend on fitting model parameters. With regard to oculomotor activity, we did not model eye movements, but used real traces recorded from human subjects during measurements of contrast sensitivity. With regard to neuronal properties, we implemented standard M and P filters obtained from the neurophysiological literature and frequently adopted by modeling studies ***(Croner and Kaplan, 1995; Benardete and Kaplan, 1997***, 1999). We chose to estimate the CSF by linearly combining M and P responses in fixed ratio, because this was the simplest model. But we note that other ways of combining M and P signals will yield very similar conclusions, since the space-time inseparability originate from the visual input rather than the neuronal models. Our two parameters (the global gain at a given temporal frequency and the ratio of M-P contributions, see Eq. 7 in the Methods section) were merely used to quantitatively align the modeled CSF with the experimental data. They have no role in explaining the shape of the CSF and its band- to low-pass transition.

In addition to providing a comprehensive explanation of the CSF, our study makes important predictions at different levels. At the neural level, our results predict that the response selectivity of RGCs will change when measured in the presence and absence of the fixational motion of the retinal image. Neurophysiological studies already suggest that fixational eye movements are an important component of visual encoding ***(Guretal., 1997; Leopold and Logothetis, 1998; Martinez-Conde et al., 2000; Ölveczky et al., 2003; Kagan et al., 2008; Meirovithz et al., 2012; Mcfarland et al., 2016).*** Eye jitter has been found to reduce redundancy in the responses of retinal neurons ***(Segal et al., 2015)*** and to synchronize them, enhancing visual features ***(Greschner et al., 2002)*** even beyond the physiological limitations imposed by photoreceptors spacing ***(¡uusola et al., 2016).*** Furthermore, retinal ganglion cells have been found that may distinguish between the global motion given by fixational eye movements and the local motion of objects ***(Ölveczky et al., 2003).*** Yet, retinal responses are traditionally measured with the eyes immobilized, a condition in which RGCs tend to exhibit relatively strong responses at low spatial frequencies ***(Croner and Kaplan, 1995).*** We predict that with normal fixational drift, because of the spatial frequency amplification in the retinal input (Fig. 2D), neuronal sensitivity to higher spatial frequencies will be enhanced and sensitivity to low spatial frequencies suppressed. As a consequence, RGCs should peak at higher spatial frequencies and exhibit a more pronounced band-pass behavior with normal fixational instability. This prediction is difficult to test in vivo, because of the need to completely stabilize the retinal input, but it can be thoroughly investigated in-vitro, where the motion of the retinal image is under full experimental control.

At the perceptual level, an interesting observation comes from the changes in the frequency content of the retinal input shown in Fig. 5 A. The amplitude of fixational instability regulates the power available in different spatial frequency bands. Specifically, the smaller the amount of retinal image motion, the more the range of amplification shifts to higher spatial frequencies. The visual system could, in principle, exploit this relationship by dynamically matching the spatial scale of eye drift to the frequency content of the visual scene, or the frequency range that is task-relevant. Within a certain range, smaller drifts would optimize information accrual when foveating on regions rich in high spatial frequencies. This effect could not only be directly driven by the stimulus in a bottom-up fashion, but also be used to meet top-down demands in high-acuity tasks. Indeed, several studies support the idea that humans can control the amount of their ocular drift ***(Steinman et al., 1973; Cherici et al., 2012; Poletti et al., 2015).*** In the same vein, the relationship between fixational drift and the frequency content of the retinal input may also explain individual perceptual differences. Subjects with relatively smaller drifts are expected to perform better in tasks in which high spatial frequencies are critical. Studies that quantitatively relate the characteristics of fixational eye drift to visual perception are needed to investigate these predictions.

Furthermore, our model predicts that manipulating temporal modulations from eye drift will affect performance. We have shown that reducing the amount of the retinal jitter well matches the overall reduction in contrast sensitivity as well as the shift to higher spatial frequencies observed in experiments of retinal stabilization. In the other direction, enlarging fixational jitter increases the amount of power available at low spatial frequencies predicting an improvement in contrast sensitivity in this range. This prediction is consistent with the improvements in word and object recognition reported in patients with central visual loss, when images or text are jittered or scrolled ***(Watson et al., 2012; Harvey and Walker, 2014; Gustafsson and Inde, 2004).*** The spatial frequency band of retinal ganglion cells decreases with eccentricity and enlarging retinal image motion has the effect of bringing more power in their range of sensitivity.

Our study also has clinical implications, as it predicts that disturbances in fixational oculomotor control will affect visual sensitivity. Oculomotor anomalies and impaired sensitivity co-occur in a variety of disorders, including conditions as diverse as dyslexia ***(Stein and Fowler, 1981,1993)*** and schizophrenia ***(Dowiasch et al., 2014; Egaña et al., 2013).*** Patients with these conditions exhibit similar visual def cits including reduced sensitivity ***(Lovegrove et al., 1980a,b; Slaghuis, 1998),*** low-level visual impairments ***(Eden et al., 1996; Li, 2002; Butler et al., 2001; Kim et al., 2006)*** and reading disabilities ***(Revheim et al., 2006)*** possibly caused by the disturbances in low-level vision ***(Revheim et al., 2006; Lovegrove et al., 1980a).*** Our results suggest a potential link between f¡ne-scale eye movements and these visual def cits, which has not yet been investigated and which may inspire novel therapeutic approaches.

## Methods

### Data collection and analysis

To examine the influences of eye movements on visual sensitivity, neuronal models were exposed to reconstructions of the input signals typically experienced by observers in experiments of contrast sensitivity. To this end, we captured the oculomotor traces recorded during contrast sensitivity measurements, in trials in which contrast was close to threshold, and used these traces to move the stimuli presented as input to the models. Methods for the collection and analysis of eye movements data as well as perceptual results have already been described in previous publications and are only briefly summarized here. See ***Mostofi et al. (2016)*** and ***Boi et al. (2017)*** for further details. This section focuses on the methods that are novel to this study.

#### Subjects

Eye movements were recorded from 5 observers (all females, age range 21-31). To optimize the precision of the recordings, only subjects with normal, uncorrected vision took part in the study. Informed consent was obtained from all participants following the procedures approved by the Boston University Charles River Campus Institutional Review Board.

#### Apparatus

Stimuli were displayed on a gamma-corrected fast-phosphor CRT monitor (Iyama HM204DT) in a dimly-illuminated room. They were observed monocularly with the left eye patched, while movements of the right eye were recorded by means of a Dual Purkinje Image eyetracker (Fourward Technology) and sampled at 1 Khz. This system has a resolution – measured by means of an artif cial eye – of approximately 1’ ***(Crane and Steele, 1985; Ko et al., 2016).*** A dental imprint bite bar and a head-rest prevented head movements. Stimuli were rendered by means of EyeRIS, a custom system that enables precise synchronization between oculomotor events and the refresh of the image on the monitor ***(Santini et al., 2007).***

#### Stimuli and Procedure

To determine the characteristics of fixational eye movements during a typical psychophysical CSF measurement, we used a standard grating-detection paradigm (see ***Mostof et al. (2016)*** for the behavioral data). In a forced-choice procedure, observers detected 2D Gabor patterns oriented at ±45°. Their contrast varied across trials following PEST ***(Taylor and Creelman, 1967).*** The frequency and standard deviation of the Gabor were 10 cycles/deg and 2.25° respectively. Stimuli were displayed over a uniform f eld with luminance of 21 ***cd/m^2^.*** Oculomotor traces were segmented in complementary periods of drift and saccades based on a speed threshold of ***2^ĉ^/s (Mostof et al., 2016).*** Only oculomotor traces collected around threshold levels of sensitivity and that contained no saccades, microsaccades or blinks were used in this study.

Modeled neurons were exposed to the same retinal input experienced by human participants, identically replicated at all spatial frequencies. Gratings were presented for 3.2 s. They were smoothly ramped up and down in contrast at the beginning and end of the trial by means of the modulating function ***M(t)*** and also modulated in time at frequency *ω_t_*, (*ω_t_*, = 0,1, 6,16, or 22Hz). The reconstructed retinal input was thus given by:

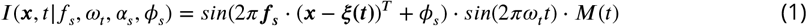

where *ξ*(***t***) = [*ξ_x_(t), ξ_y_(t)*] represents eye movements and ***f***_*s*_ = [*f_s_cos*(*α_s_*), *f_s_sin(α_s_)*] the stimulus frequency (0.1-60 cycles/deg). The orientation *α_s_* and the phase *ϕ_s_* uniformly spanned the range [0 2*π*).

#### Neural models

The mean instantaneous rate of retinal ganglion cells (RGCs) were simulated by means of standard space-time separable linear filters with transfer function:

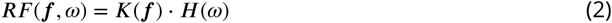

where ***f*** indicates spatial frequencies and ***ω*** temporal frequency. The spatial kernel ***K(f)*** was modeled as in ***Croner and Kaplan (1995)*** with a standard difference of Gaussians:

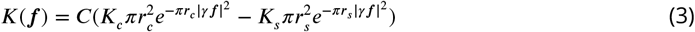

Parameters were adjusted based on the neurophysiological recordings from macaques (Table 1 in ***Croner and Kaplan (1995))*** with the scaling factor ***γ*** set to 0.5 to model the smaller receptive fields of the fovea following the magnification factor (formula 8 in ***Van Essen et al., 1984).***

The temporal kernel consisted of a series of low-pass filters and a high-pass stage ***(Victor, 1987***), to yield a transfer function ***H(ω)***:

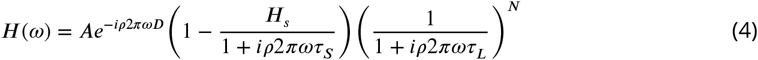

Parameters were adjusted on the basis of neurophysiological responses (M cells: median values in Table 2 in ***Benardete and Kaplan, 1999;*** P cells: median values in Table 2 in ***Benardete and Kaplan, 1997)*** with the scaling factor ***<*** set to 1/1.6 to include the effects of large stimuli on retinal responses (Fig. 7B in ***Alitto and Usrey, 2015).***

#### Estimating contrast sensitivity

The main assumption of our model is that the visual system is insensitive to temporal stimulation at 0Hz so that spatial sensitivity is entirely driven by temporal transients. For this reason, we estimated the predicted CSF on the basis of cell responses to input changes.

For each spatial frequency ***f_s_*** of the grating, we first estimated the space-time power spectrum of the retinal input ***P_I_*(*f, ω*)** by averaging the square of the absolute value of the Fourier transform of Eq. 1 across trials, stimulus’ orientations ***α_s_*** and phases ***φ_s_***. Since both ***P_I_***(*f, ω*) and the spatial kernels ***K(f)*** possess circular symmetry in spatial frequency, we reduced the spatial dimensionality from 2D to 1D by radial averaging. We then computed the power spectrum of neuronal responses ***O(f,ω)*** by multiplying the space-time power spectrum of the retinal input ***P_I_*(*f,ω*)** by the transfer functions of the cells’ filters:

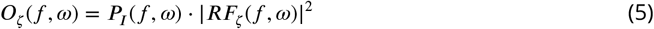

where ***RF_ξ_*(*f, ω*)**, with ***ζ = M*** or ***P***, represents the Fourier transform of M or P cells’ receptive fields (Eq.2).

Finally, we evaluated the CSF at each spatial frequency ***f***, by computing the square root of the integrated temporal power across all non-zero temporal frequencies:

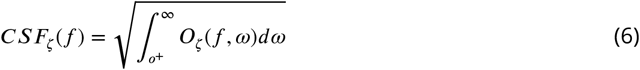

where ***O_ζ_*** represents the power spectrum of M or P responses. The integral in Eq. 6 was computed numerically. To avoid artifacts from finite bandwidth, the first two temporal samples of the spectrum were discarded so that integral over temporal frequency started from ***ω*** = 0.63*Hz*.

The predicted CSF was then estimated, for each condition, by a linear combination of the contrast sensitivities of the two types of neurons, ***CSF_M_(f)*** and ***CSF_P_(f)***:

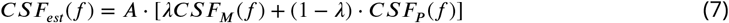

where *λ* (*λ* = 0.57 for all conditions) weighs the contributions of the M and P populations and *A* is a global rescaling coefficient.

Note that the parameters *A* and *λ* were merely used to quantitatively align model predictions with classical data, but had no role in explaining our findings. That is, the emergence of a space-time inseparability in the CSF, was neither caused by the specif c value of *λ* (both M and P cells show this transition; Fig. S2A-B)) nor by the global scaling factor A, which had no effect on the shape of the predicted CSF. We chose to linearly combine the contributions of M and P neurons because this was the simplest model. However, use of other models (e.g., the maximum of either population at each spatial frequency *f*) produced virtually the same results given the robustness of the underlying phenomenon.

The same procedure was used to estimate the CSF in the case of no eye movements and retinal stabilization (Figs. 3, 4 and 5). In the former condition (no eye movements), *ξ(t)* was set to zero in Eq. 1. In the latter condition (retinal stabilization), we modeled the retinal image motion by means of a 2D random walk process, but with reduced diffusion coefficient (*D*=2 rather than the normal value D=250). Brownian motion, with *D* in the range 100-350, is known to be a good model for the normal retinal image motion when the head is not immobilized ***(Aytekin et al., 2014).***

**Figure 5.**
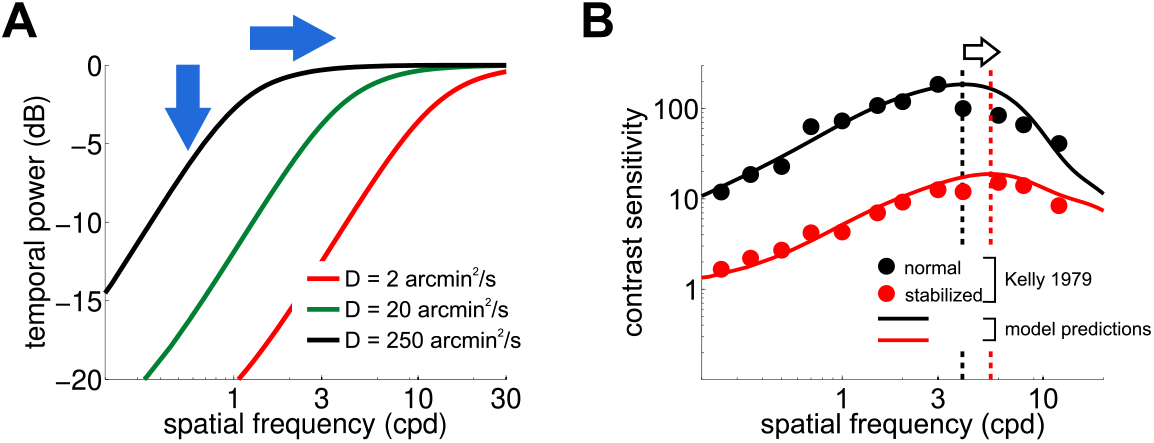
Consequences of retinal stabilization. (A) Spatial spectral density of the luminance modulations resulting from a Brownian model of retinal image motion with different diffusion constants. Lowering *D* both attenuates the power available at each spatial frequency (vertical arrow) and shifts the distribution to higher spatial frequencies (horizontal arrow). (B) Predicted contrast sensitivity under retinal stabilization. Sensitivity is reduced and shifted to higher spatial frequencies. Dashed vertical lines mark the maxima of the two curves (color coded according to their *D* in panel A). Results quantitatively match classical experimental data from ***Kelly (1979).*** CSFs predicted separately from the responses of M and P neurons are shown in Fig. S2 C.

**Figure S1.**
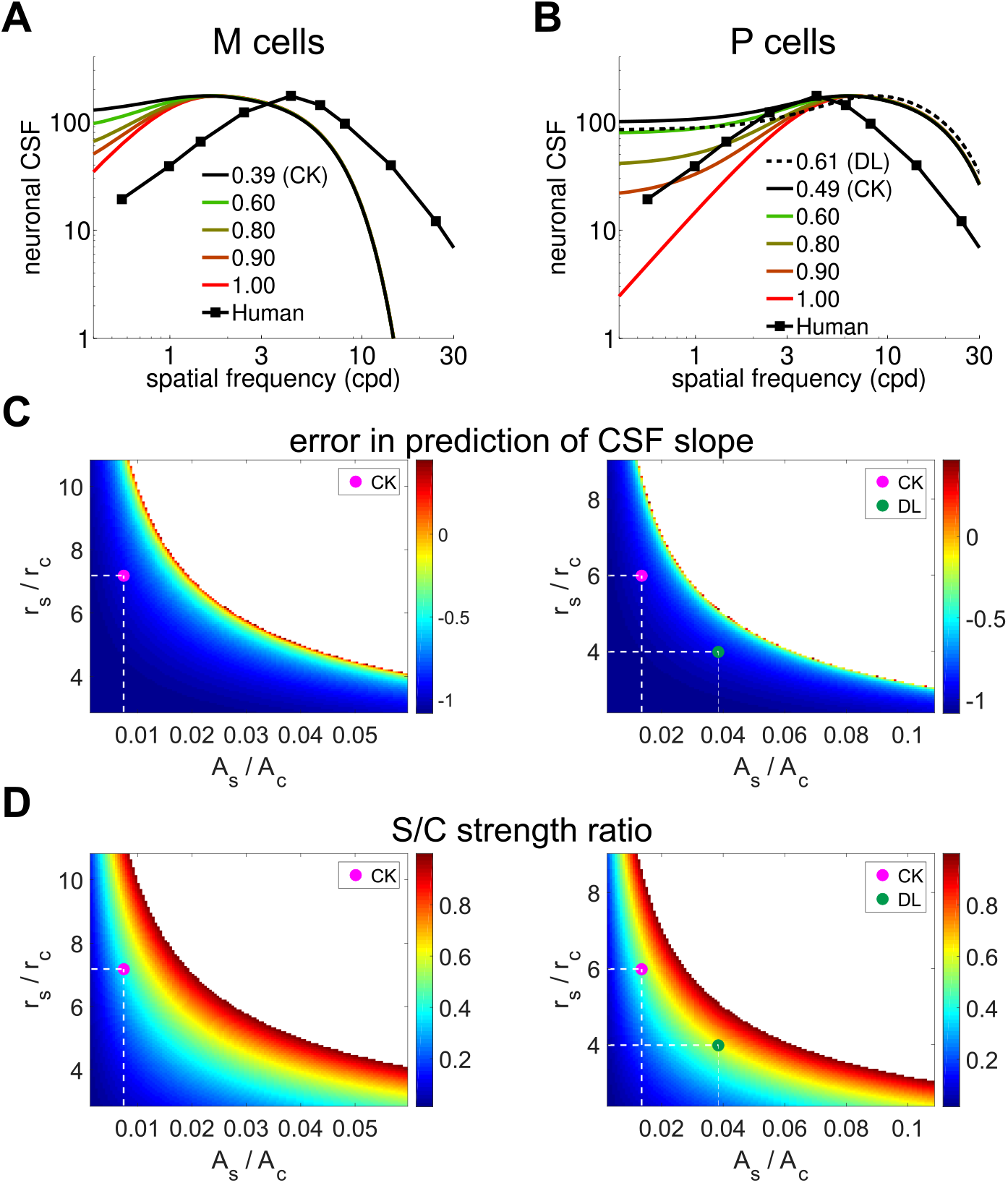
(A, B) Spatial sensitivity in standard models of magno-(A) and parvo-cellular cells (B) as a function of the ratio between the strengths of their center and surround. The human CSF from Fig. 1A is also plotted for comparison. ‘DL’ and ‘CK’ label the ratios measured experimentally by ***Derrington and Lennie (1984)*** and ***Croner and Kaplan (1995)*** respectively, from the medians of their reported values. All other parameters were set as described in the Methods section. (C, D) Full parametric analysis of the difference in slope at low spatial frequencies between the human CSF and the spatial sensitivity of difference-of-Gaussians models. Each point in the map shows the slope deviation resulting from a particular ratio between surround and center amplitudes (***A_s_/A_C_***, horizontal axis) and between radii (***r_s_/r_c_***, vertical axis) in the models (Eq. 3). A value of zero represents perfect matching between the CSF and the receptive fields profile;negative/positive values indicate that the neuronal filter is less/more attenuated than the CSF. Values of the parameters for which the slope could not be computed because the receptive field did not exhibit a band-pass behavior are indicated by white. The magenta and greed dots mark parameters measured experimentally by ***Croner and Kaplan (1995)*** and ***Derrington and Lennie (1984)*** respectively (dashed lines). (D) Ratio between center/surround excitation and inhibition. A value of 1 indicates that center and surround have the same strength. Legends and symbols are as in C. Comparison of panels C and D shows that a slope similar to that of the human CSF can only be obtained for close balance between excitation and inhibition. These values differ greatly from those measured experimentally (magenta dot).

**Figure S2.**
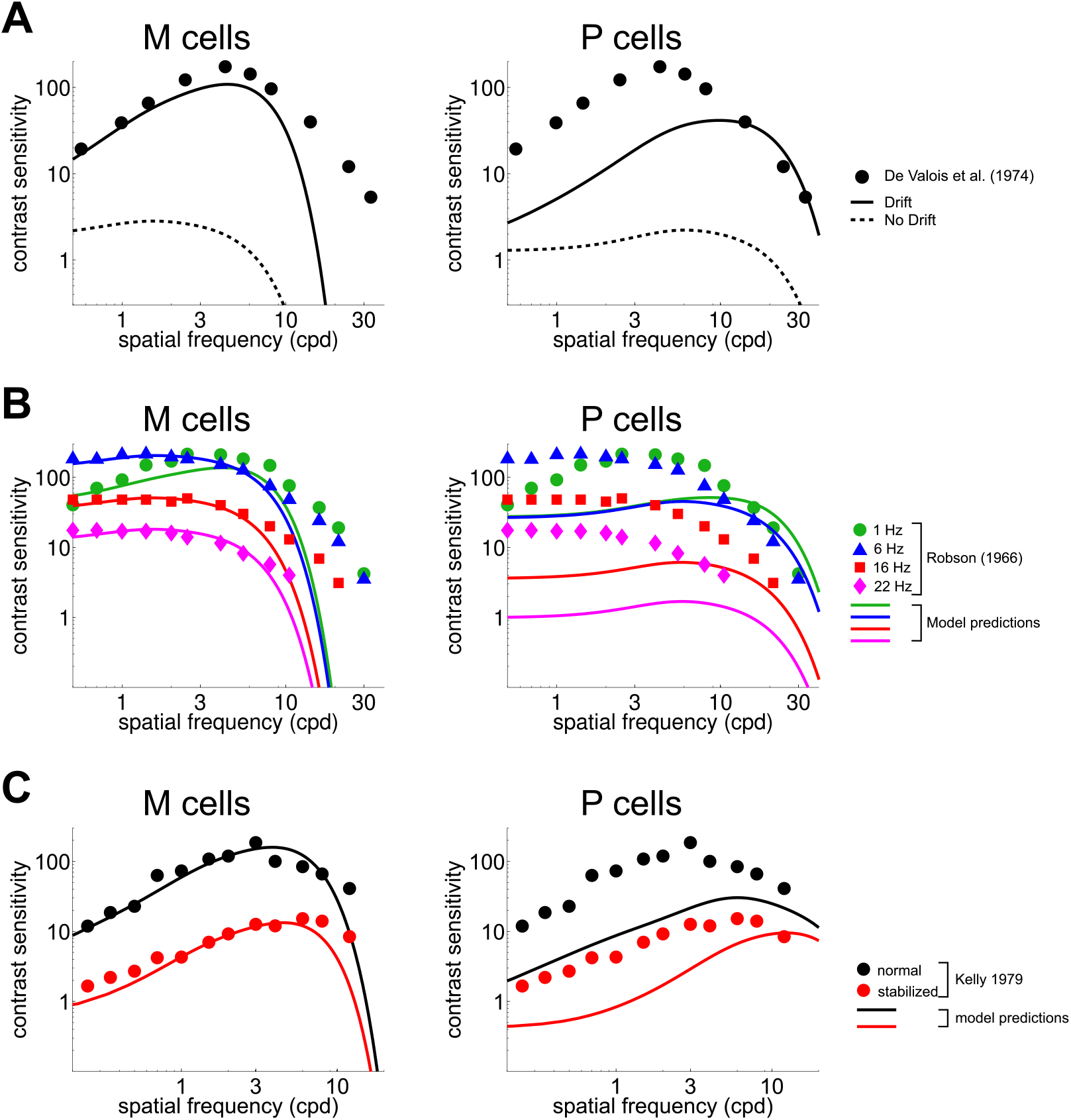
CSF predicted separately from the responses of M and P cells. Legends and symbols in panels A, B and C are as in Fig. 3, Fig. 4B and Fig. 5B, respectively.

